# Subtype-specific differences in susceptibility to monoclonal antibodies and vaccines among contemporary RSV-A and RSV-B isolates

**DOI:** 10.64898/2026.06.09.731176

**Authors:** Paula Eillanny Silva Marinho, Dibya Ghimire, João Tsai Silva, Jamie Devlin, Sunyoung Kwon, Gordon Adams, Patricia Jorquera, Jeremy Luban, Jacob E. Lemieux

**Affiliations:** Massachusetts General Hospital, Boston, MA; Moderna Tx; University of Massachusetts Chan Medical School, Worcester, MA

**Keywords:** Respiratory syncytial virus, RSV subtypes, Monoclonal antibodies, Antigenic divergence, Neutralization

## Abstract

Respiratory syncytial virus (RSV) continues to circulate at high levels despite the introduction of new monoclonal antibodies (mAbs) and vaccines targeting the prefusion F (pre-F) protein. We analyzed the viral genome sequences of 133 RSV clinical samples collected during the 2022–2023 and 2023–2024 seasons, and selected representative isolates for phenotypic testing. We selected four RSV A and four RSV B replication-competent, sequence-verified stocks that were assessed for replication kinetics *in vitro* and neutralization sensitivity by a panel of F-targeting mAbs and polyclonal sera using a rabbit vaccination model. Isolates retained sensitivity to all mAbs tested. We detected no mutations in mAb binding sites in the isolates tested. RSV-A isolates were slightly less susceptible to multiple mAbs than RSV-B isolates. Antigenic cartography revealed a separation of antibody responses by subtype: RSV-A isolates clustered together and aligned with lower neutralization by antibodies such as MPE8 and 101F, whereas RSV-B isolates formed a distinct cluster associated with higher mAb susceptibility. In a rabbit vaccination model, RSV-A–only sera efficiently neutralized all RSV-A isolates and the RSV-B1 reference strain but showed diminished activity against contemporary RSV-B isolates. RSV-B–only sera displayed balanced neutralization across both subtypes. Combined RSV-A+B immunization produced uniformly strong responses to all isolates, suggesting that multivalent exposure may overcome subtype-specific antibody polarization. Collectively, our results demonstrate consistent antigenic divergence of RSV subtypes and underscore the importance of considering genetic and phenotypic divergence for F-directed immunoprophylaxis.

**Importance:** RSV remains a major cause of lower respiratory tract infections. Although new mAbs and vaccines have been approved in recent years, the impact of circulating genetic diversity on these therapeutics is incompletely understood. In this study, we demonstrate how recent RSV isolates respond to a panel of mAbs and whether sera from rabbits vaccinated with mRNA vaccines effectively neutralize these isolates. We reveal that RSV-A isolates are more resistant to neutralization by mAbs than RSV-B isolates, and the bivalent vaccine elicits broader neutralizing responses than the monovalent vaccine. This study provides insights into how viral diversity may influence antibody-mediated protection and suggests that recent RSV-B isolates may respond differently from the laboratory-adapted strains commonly used in research.

## Introduction

Respiratory Syncytial Virus (RSV) is one of the leading causes of lower respiratory tract infections, including pneumonia and bronchiolitis worldwide ^1–3^. Globally, RSV is estimated to affect around 64 million people and 160,000 deaths ^4^. RSV season occurs yearly between October and April in the United States. In the 2024-2025 season in the United States alone, RSV was responsible for an estimated 190,000-350,000 hospitalizations, and approximately 23,000 deaths ^5^. RSV remains a major cause of lower respiratory tract infections, imposing a substantial burden on geriatric and infant age groups and hospitalization rates ^6,7^.

RSV is a negative-sense single-stranded RNA virus that belongs to the genus *Orthopneumovirus.* It contains a non-segmented genome that codes 11 proteins. RSV is subdivided into two subtypes, A and B, which are characterized by antigenic drift in the G, M, and N genes^8^. Two surface glycoproteins, the Fusion glycoprotein (F) and attachment protein (G) are the main antigenic determinants for neutralizing antibodies ^9^. The F protein is highly conserved between RSV subtypes A and B, with approximately 90% sequence identity ^9^, and the majority of neutralizing antibodies targeting this protein exhibit cross-reactive activity against both subtypes ^10^. The RSV F protein is a trimeric class I viral fusion glycoprotein that plays a central role in mediating viral entry through membrane fusion with host airway epithelial cells. F has been widely utilized as a primary antigenic target for vaccines and monoclonal antibodies. It is synthesized as an inactive precursor (F0) that is cleaved by host furin-like proteases into the F1 and F2 subunits, priming the protein for fusion. The F protein exists in a metastable prefusion (preF) conformation that undergoes an irreversible structural rearrangement to the highly stable postfusion (postF) state during the fusion process.

In 2013^11^, the successful stabilization and recombinant expression of the preF conformation represented a major advance, enabling the rational design of RSV vaccines and monoclonal antibodies^12^. Structurally, the preF form displays multiple key neutralizing epitopes, including antigenic sites Ø, I, II, III, IV, and V, whereas the postF conformation retains only a subset of these sites, primarily I, II, and IV. Antibodies targeting prefusion conformation, specifically antigenic sites Ø and V, have been shown to exhibit markedly higher neutralization potency. Antibodies targeting the postfusion conformation are mostly non-neutralizing ^13^.

Therapeutic options for RSV infection remain limited, highlighting the importance of prophylaxis. In the United States, RSV prevention is primarily achieved through passive and active immunization approaches targeting populations at highest risk of severe disease. Two vaccines, Arexvy (GlaxoSmithKline) and Abrysvo (Pfizer), both based on the pre-F protein, were approved by the U.S. Food and Drug Administration (FDA) in 2023 for the prevention of lower respiratory tract disease caused by RSV in adults 60 years of age and older ^14,15^. Abrysvo is also approved for use in pregnant women at 32–36 weeks’ gestation to protect infants from birth through 6 months of age^15^. In 2024, the FDA approved MRESVIA (Moderna’s mRNA-1345), an mRNA vaccine encoding the stabilized pre-F protein, also for adults aged 60 years and older ^16^. Despite these advances, no RSV vaccine has yet been approved for direct use in infants.

Prophylaxis with mAbs remains the primary pharmacologic strategy for preventing severe RSV disease in infants. For many years, palivizumab (Synagis®), a mAb targeting antigenic site II of the RSV F protein, was the only approved option and was limited to high-risk pediatric populations ^17^. More recently, broader preventive approaches have emerged. In July 2023, the U.S. FDA approved nirsevimab (Beyfortus®), a long-acting mAb directed against antigenic site Ø of preF, for use in all infants entering their first RSV season and in selected high-risk children entering their second season ^18,19^. Subsequently, in 2025, clesrovimab (Enflonsia®), which targets antigenic site IV of the F protein, was also approved for prophylactic use in neonates and infants at risk of RSV infection^20,21^. Together, these approvals expand the options for passive immunization against RSV in early life, with each antibody targeting distinct antigenic sites on the F protein ^20^.

Genomic surveillance for RSV is relatively limited compared to other common respiratory viruses such as Influenza and SARS-CoV-2, but surveys of viral diversity have demonstrated that multiple RSV A and B clades are extant^22–24^, but the impact of circulating genetic diversity on monoclonals and vaccines is incompletely understood. To address these gaps, we isolated RSV strains from nasal swabs collected at Massachusetts General Hospital between November 2022 and March 2024 and characterized their virologic properties *in vitro*. We assessed the extent of immune escape of these isolates against clinically approved mAbs, as well as several preclinical antibody candidates.

## Results

### Isolation of RSV from Human Clinical Samples

To isolate a panel of RSV isolates, we inoculated 133 nasal swabs from 113 unique patients and obtained CPE in 41 specimens. The number of RSV isolates obtained varied between subtypes (Figure S1). During the 2022–2023 and 2023–2024 seasons, we recovered 32 isolates, 31 RSV-A and 1 RSV-B isolates, from 77 nasal swabs. In the following season (2023–2024), we obtained 9 isolates, including 3 RSV-A and 6 RSV-B, from 56 nasal swabs. We selected eight of these isolates, four RSV-A and four RSV-B, from phylogenetically representative genotypes, for further analysis (Table 1 and Figure 1).

**TABLE 1.** RSV strains isolated and used in this study.

| Name | Subtype | Season | Strain Designation | Genbank accession number | F protein AA substitutions |
| --- | --- | --- | --- | --- | --- |
| MA-MGB-RSV-0016 | RSV-B | 2023-2024 | MA-Broad_MGH-16016/2023 | PV178377.1 | F_T529A |
| MA-MGB-RSV-0030 | RSV-B | 2023-2024 | MA-Broad_MGH-16030 | PV260458.1 | F_T529V |
| MA-MGB-RSV-0089 | RSV-B | 2023-2024 | MA-Broad_MGH-16089/2023 | PV178453.1 | F_I11V<br>F_R42K<br>F_T529A |
| MA-MGB-RSV-0112 | RSV-B | 2022-2023 | MA-Broad_MGB-13686 | OQ024160.1 | R42K<br>F_T529A |
| MA-MGB-RSV-0115 | RSV-A | 2022-2023 | MA-Broad_MGB-13689 | OQ024139 | F_T103A<br>F_A122T |
| MA-MGB-RSV-0119 | RSV-A | 2022-2023 | MA-Broad_MGB-13693 | OQ024156 |  |
| MA-MGB-RSV-0151 | RSV-A | 2022-2023 | MA-Broad_MGB-13725 | OQ024143 | F-T12I<br>F_M115I<br>F_A122T |
| MA-MGB-RSV-0152 | RSV-A | 2022-2023 | MA-Broad_MGB-13726 | OQ024141 | F-T12I<br>F_M115T<br>F_A122T |
| RSV-A2 | RSV-A |  | hRSV/A/Australia/A2/1961 | KT992094.1 |  |
| RSV-B1 | RSV-B |  | RSV B1 wt | AF013254.1 |  |

**Figure 1.**
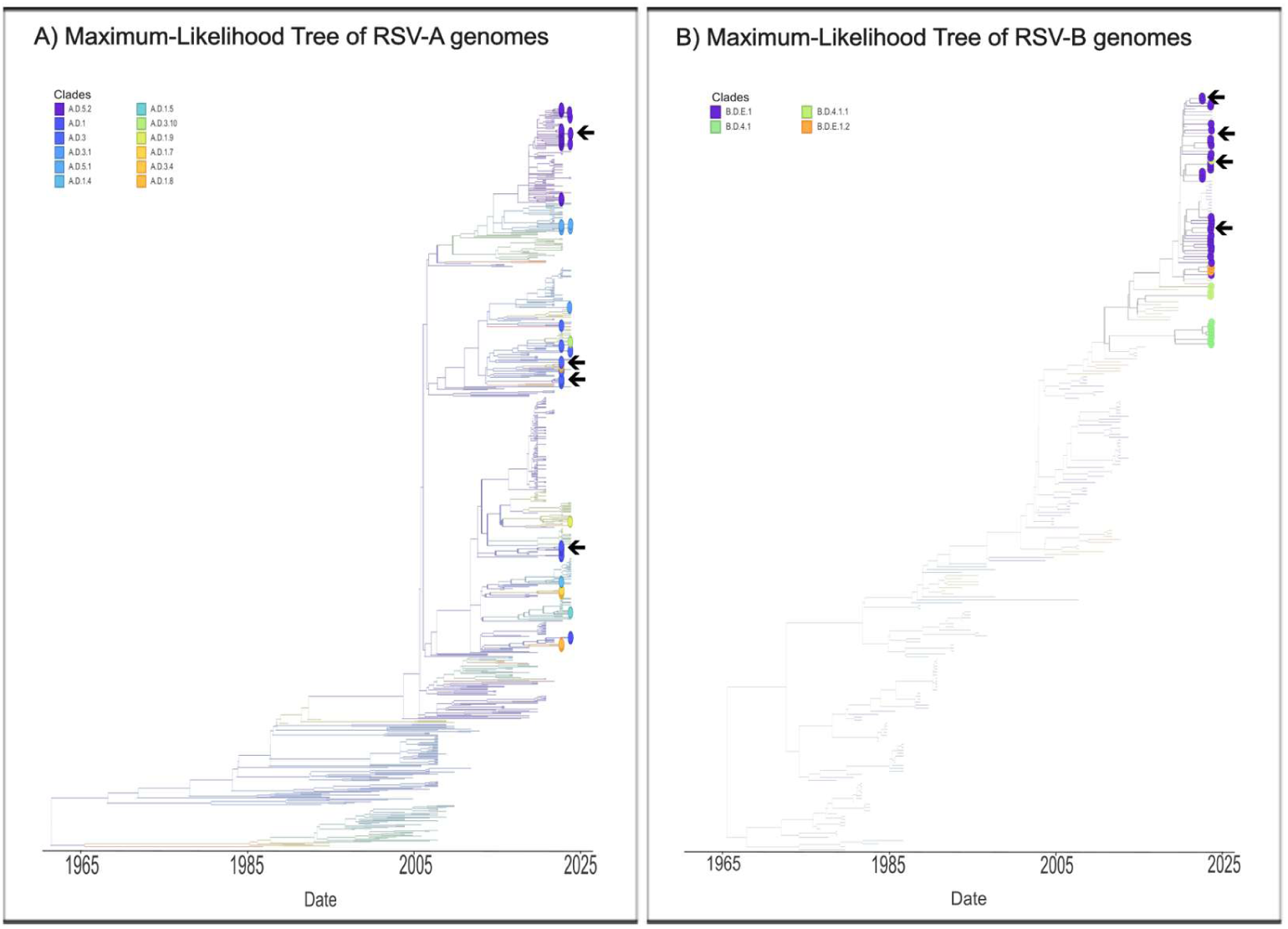
Phylogenetic analysis of RSV genomes. A) RSV-A sequences from 55 attempted isolates. B) RSV-B sequences from 46 attempted isolates. Samples selected for downstream analyses are indicated by arrows.

### Phylogenetic Analysis and Genetic Divergence

We performed whole genome sequencing on viral RNA obtained nasal swabs from all 133 samples from 113 unique patients. Phylogenetic trees for RSV-A and RSV-B sequences derived from clinical samples are shown in **Figure 1**. RSV-A sequences clustered within the clade A.D.1, A.D.3 and A.D.5, consistent with the circulation of contemporary RSV-A lineages^22–24^. RSV-B sequences clustered within the clade B.D.4.1, B.D.4.1.1, B.D.E.1 and B.D.E.1.2, also aligning with globally prevalent RSV-B strains during the study period. We resequenced the isolates in Table 1 and confirmed that the genome sequences of the isolated viruses were identical to those obtained directly from the corresponding patient samples, indicating no adaptive changes during isolation.

We next analyzed intra-group genetic diversity among RSV clinical isolates. Nucleotide divergence relative to clinical consensus sequences was significantly higher in RSV-A compared to RSV-B for both genes (Figure 2), with RSV-A isolates showing greater sequence variability within their subtype (F gene: Mann-Whitney U =1713.5, p = 1.9641× 10⁻^4^; G gene: Mann-Whitney U = 1927, p =7.7457 ×10⁻⁵).To visualize the distribution of genetic variation within a phylogenetic context, we analyzed position-specific mutations in the F and G genes for RSV-A and RSV-B. Nucleotide-level mutation analyses are presented as supplementary figures (Figures S2–S5).Amino acid variation across the F protein was mapped and epitope regions targeted by the mAbs evaluated in this study were annotated for reference (Figures 3 and 4).

**Figure 2.**
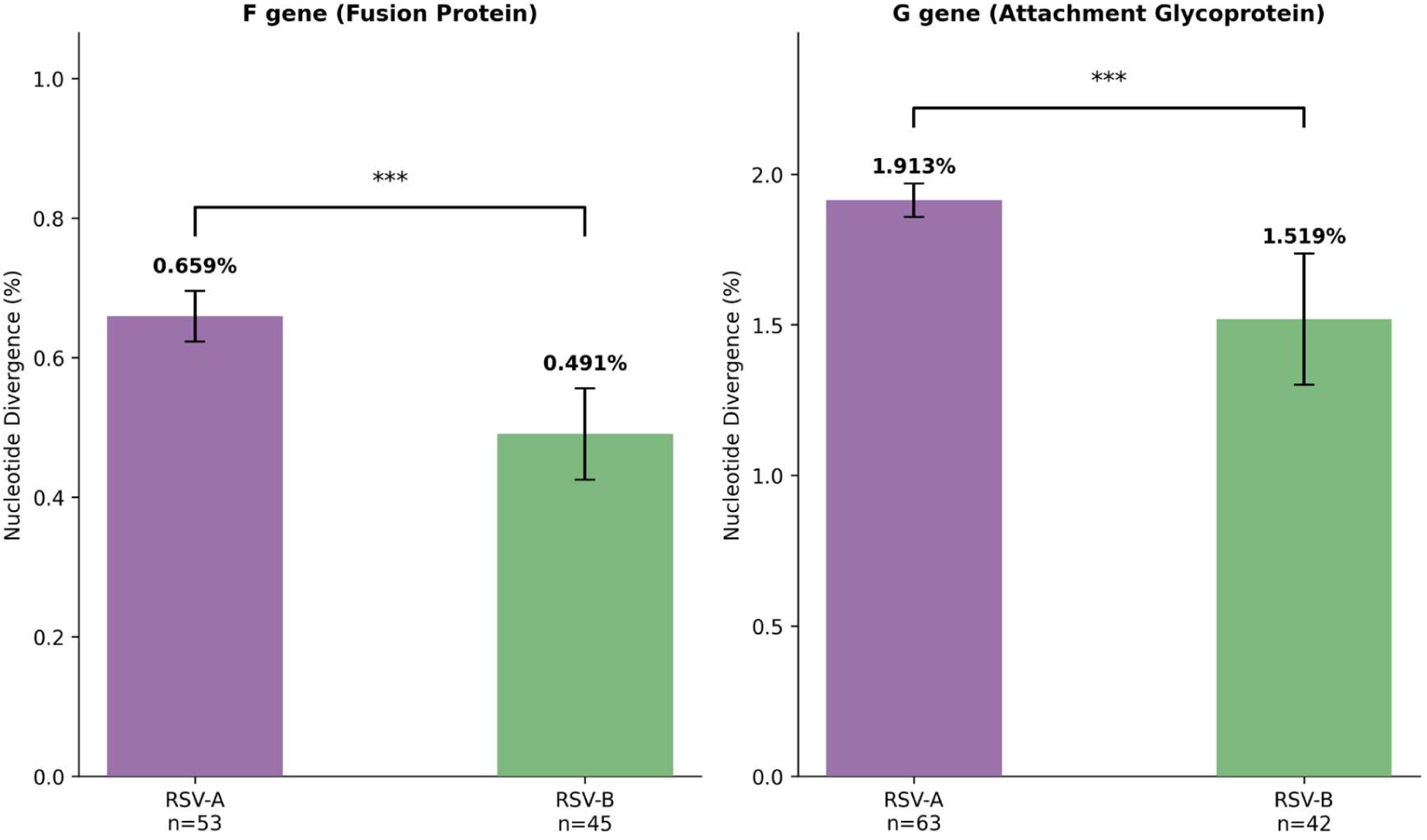
Genetic divergence within circulating RSV clinical isolates. Percent nucleotide divergence of Massachusetts (2022-2024) clinical isolates relative to their respective clinical group consensus sequences. RSV-A (purple) and RSV-B (green) are shown for F gene (A) and G gene (B). Values are shown as mean ± SEM with sample sizes indicated. RSV-A exhibited significantly higher intra-group genetic diversity than RSV-B for both genes (Mann-Whitney U test, p < 0.001 for both comparisons).

**Figure 3.**
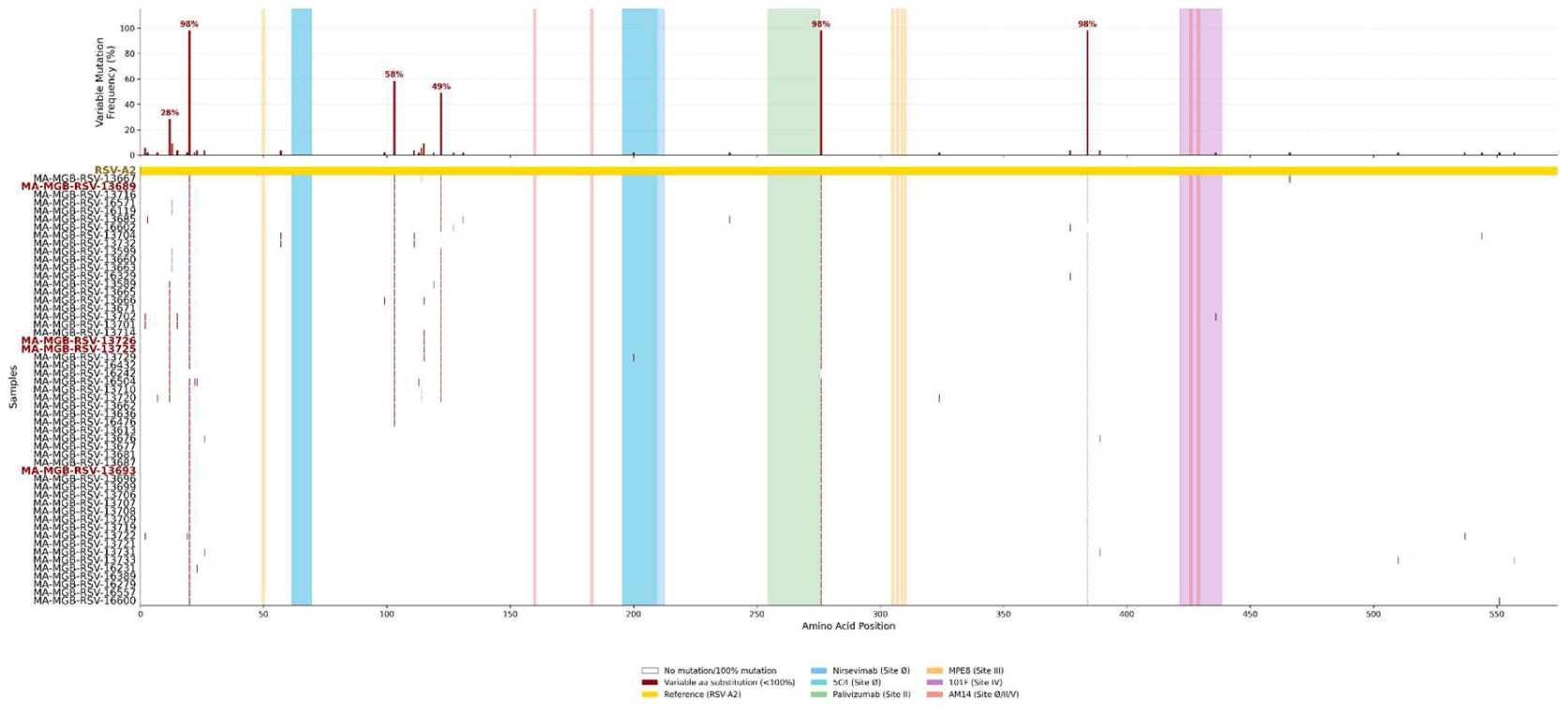
RSV-A F protein amino acid variation and monoclonal antibody epitope mapping. Phylogenetically ordered amino acid substitution analysis of RSV-A F protein (574 aa; n=53 after gene-specific quality filtering). Top panel: frequency of variable amino acid substitutions across samples relative to reference RSV-A2 (KT992094.1). Bottom panel: amino acid substitution matrix with samples ordered by phylogenetic clades (A.D.1, A.D.3, A.D.5.1, A.D.5.2 and subclades). Red labels: isolates of interest (MA-MGB-RSV-13689, MA-MGB-RSV-13693, MA-MGB-RSV-13725, MA-MGB-RSV-13726). Gold line: reference sequence (RSV-A2). White: no substitution/fixed substitution (100%). Dark red: variable amino acid substitution (<100%). Colored regions indicate epitope sites of mAbs evaluated in this study: D25 (Site Ø, blue), 5C4 (Site Ø, cyan), Palivizumab (Site II, green), MPE8 (Site III, orange), 101F (Site IV, purple), AM14 (Site Ø/II/V, red).

**Figure 4.**
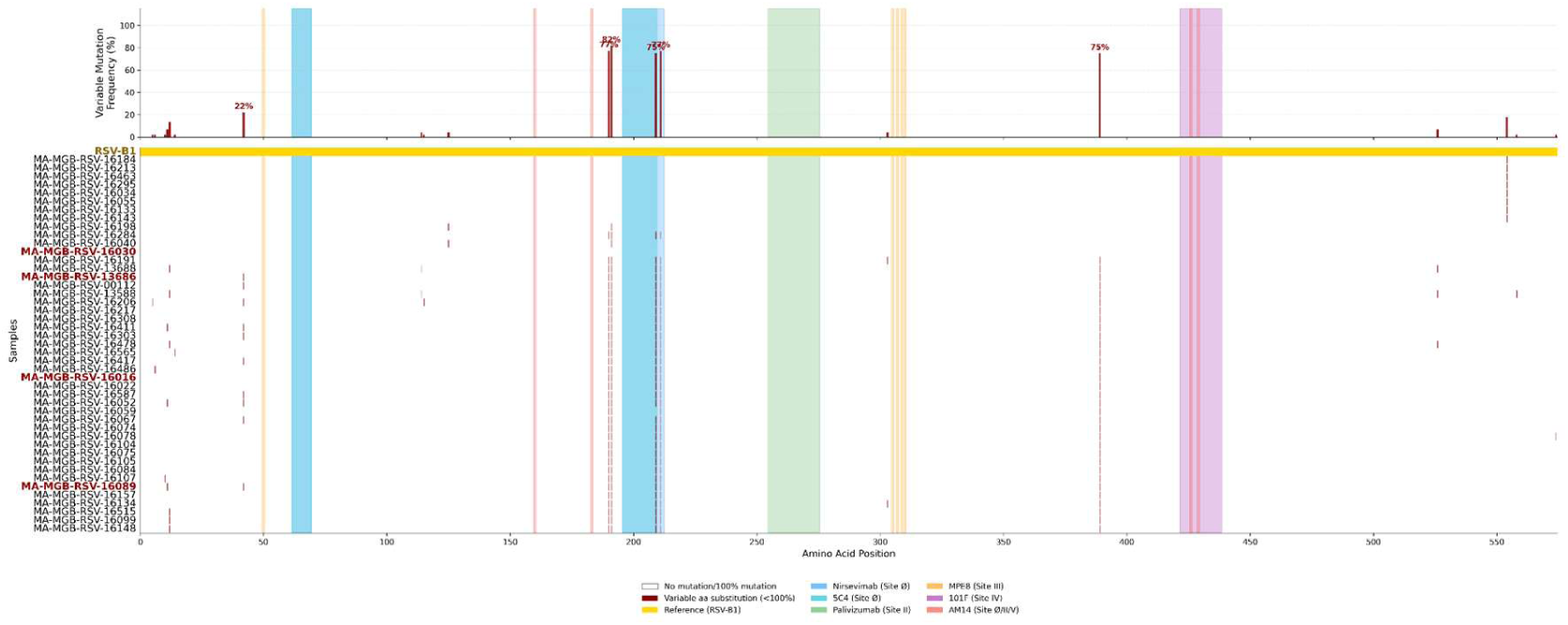
RSV-B F protein amino acid variation and monoclonal antibody epitope mapping. Phylogenetically ordered amino acid substitution analysis of RSV-B F protein (574 aa; n=45 after gene-specific quality filtering). Top panel: frequency of variable amino acid substitutions across samples relative to reference RSV-B1 (AF013254.1). Bottom panel: amino acid substitution matrix with samples ordered by phylogenetic clades (B.D.4.1, B.D.4.1.1, B.D.E.1, B.D.E.1.2). Red labels: isolates of interest (MA-MGB-RSV-16030, MA-MGB-RSV-13686, MA-MGB-RSV-16016, MA-MGB-RSV-16089). Gold line: reference sequence (RSV-B1). White: no substitution/fixed substitution (100%). Dark red: variable amino acid substitution (<100%). Colored regions indicate epitope sites of mAbs evaluated in this study: D25 (Site Ø, blue), 5C4 (Site Ø, cyan), Palivizumab (Site II, green), MPE8 (Site III, orange), 101F (Site IV, purple), AM14 (Site Ø/II/V, red).

### Viral replication kinetics

To assess viral replication, we measured growth in Hep-2 cells at 24, 48, 72, and 96 hours post-infection (hpi). All RSV-A isolates and B isolates established productive infection in HEp-2 cells (Figure 5a), with viral titers increasing over time. Both RSV-A and RSV-B replicated efficiently in HEp-2 cells. Whereas some individual isolates achieving higher titers than others, there was no significant difference between the subtypes at all timepoints except at 96 h. These differences may reflect isolate-specific variations in viral fitness or host cell interactions. Compared to historical reference strains, RSV-A2 and RSV-B1, contemporary isolates demonstrated comparable growth in HEp-2 cells, except for RSV-A2 at 24 hpi, which replicated faster initially (p < 0.01), possibly reflecting its laboratory-adapted state. To exclude a HEp-2-specific phenotype, we also tested growth in A549 cells (Figure 5b). In A549 cells, both subtypes replicated with similar kinetics, though overall titers were slightly lower than in HEp-2 cells.

**Figure 5.**
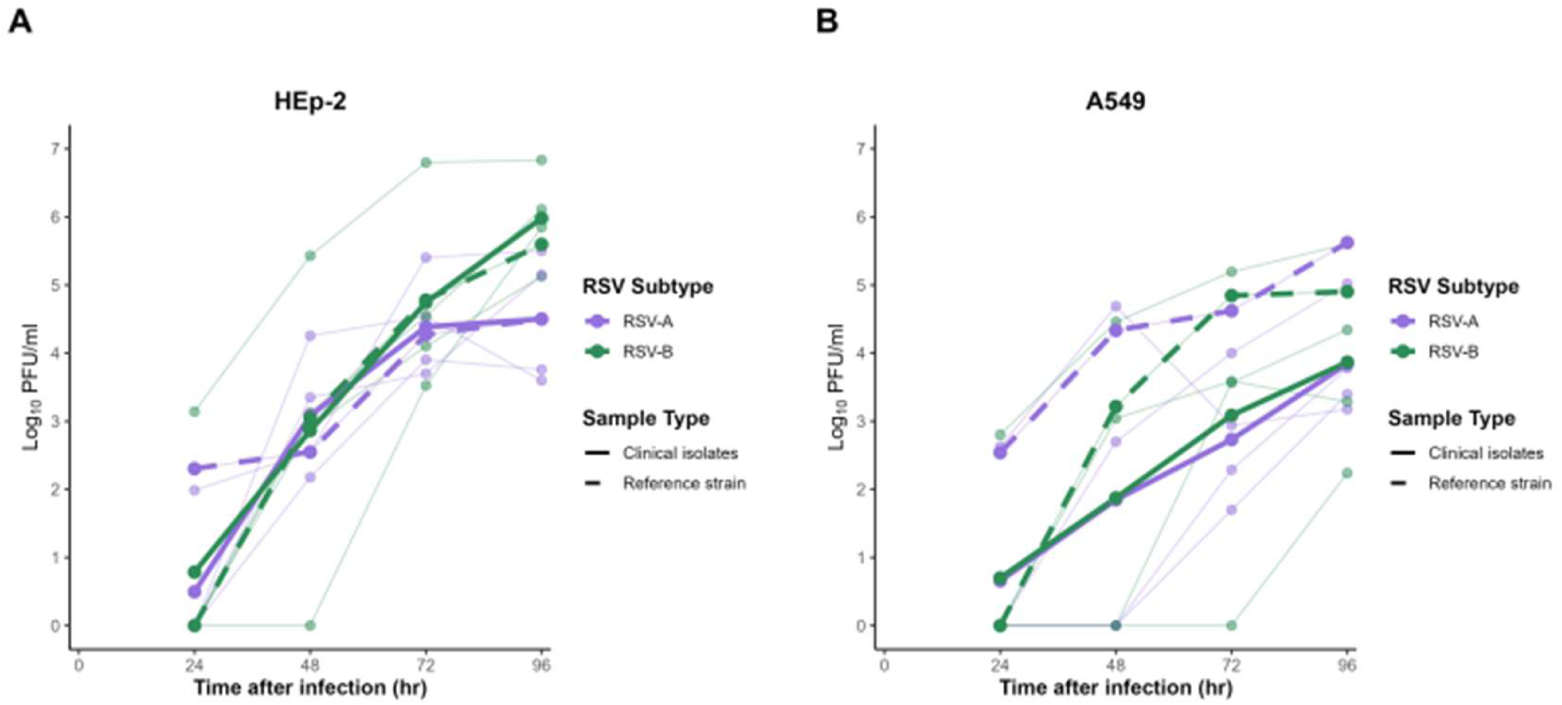
In vitro growth kinetics of RSV-A and RSV-B strains in epithelial cell lines. HEp-2 (A) and A549 (B) cells were infected with clinical isolates of RSV-A (n=4) and RSV-B (n=4) at a multiplicity of infection (MOI) of 0.01. Viral titers in cell culture supernatants were measured at the indicated timepoints post-infection by plaque assay. Each point represents the mean titer for an individual isolate from two independent experiments. Solid thicker lines represent the mean of clinical isolates; dashed lines indicate reference strains (RSV-A2 and RSV-B1). Statistical significance was determined by Student’s t-test comparing RSV-A versus RSV-B at each timepoint (**p < 0.01).

Similar to the behavior in HEp-2 cells, RSV-A2 exhibited significantly enhanced replication compared to clinical RSV-A isolates across multiple timepoints (24-96 hpi, p < 0.001). RSV-B1 showed growth comparable to clinical RSV-B isolates. Overall, RSV-A and RSV-B clinical isolates exhibited comparable viral replication in HEp-2 and A549 cells (Figure S6) with slightly higher titers observed in HEp-2 at late timepoints (72h for RSV-A, 96h for RSV-B; p < 0.01).

### Virus neutralization by RSV F-specific monoclonal antibodies

To assess antigenic characteristics of the isolates, we assessed their susceptibility to mAbs. We performed neutralization assays using a panel of antibodies targeting distinct antigenic sites on the RSV F protein (Figure 6). The panel included mAbs in clinics (nirsevimab and palivizumab) and other mAbs in preclinical studies (MPE8, AM14, 101F, and 5C4), each recognizing different structural or conformational epitopes on the F protein. (Table S1). We developed a focus-reduction neutralization test (FRNT), which could be performed in 96-well format, and confirmed its correlation with PRNT (R = 0.793; Figure S7). We obtained IC₅₀ values using FRNT for each isolate–antibody combination (Table 2). All antibodies retained neutralizing activity, although there was notable variability across isolates. Nirsevimab consistently exhibited the highest potency, with IC₅₀ values below 20 ng/mL for several isolates, including RSV-030, RSV-089, RSV-0112, RSV-0151, and RSV-A2. In contrast, palivizumab showed substantially higher IC₅₀ values, typically above 500 ng/mL, except for RSV-0151 and RSV-B1.

**FIGURE 6.**
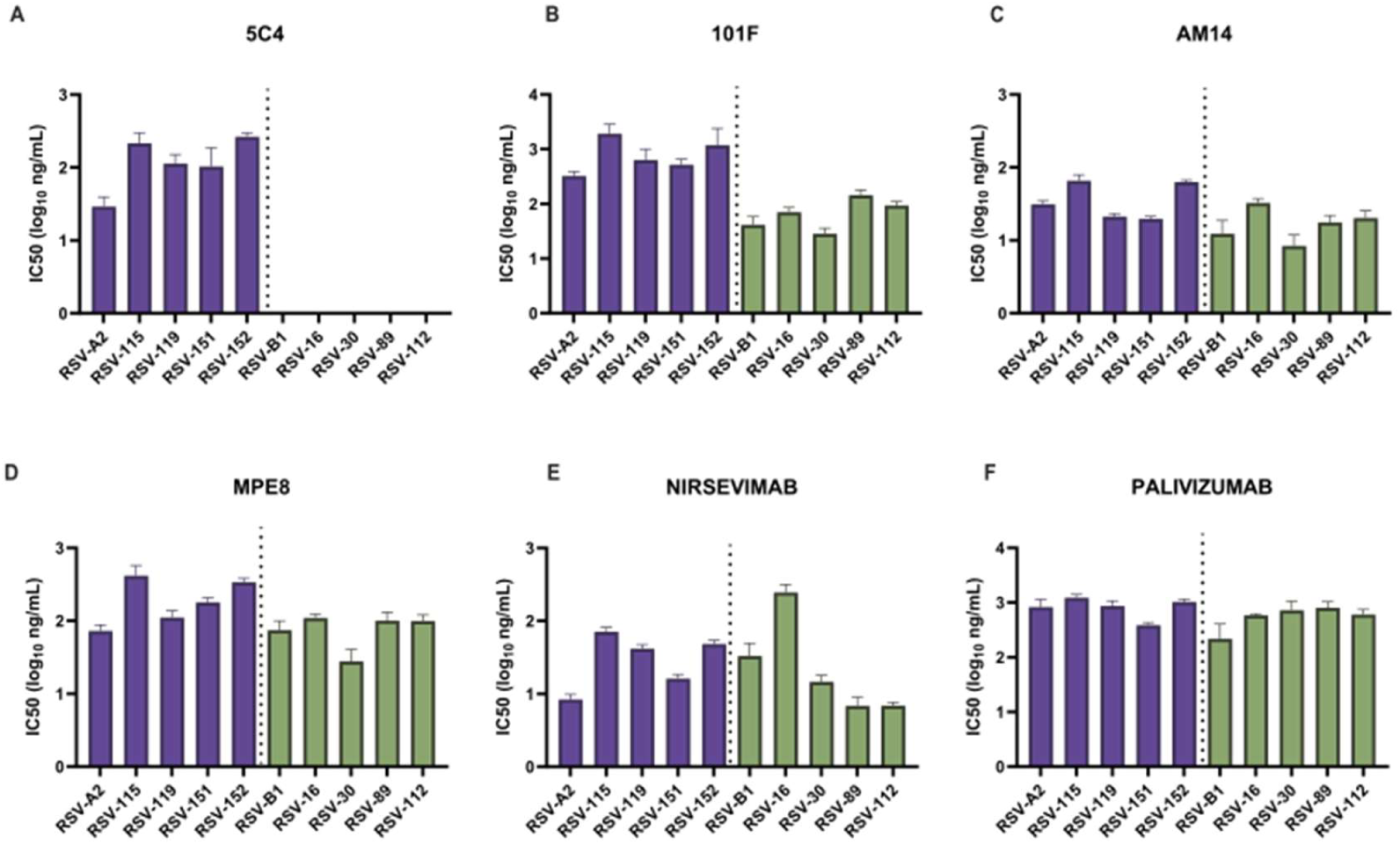
Neutralization activity of mAbs against RSV-A (purple) and RSV-B (green). IC₅₀ values were determined by nonlinear regression (log concentration vs. response variable slope), in a focus-forming assay. Statistical analysis using a two-tailed Mann–Whitney U test showed significant differences between RSV-A and RSV-B for mAbs 101F (p = 0.0079) and MPE8 (p = 0.007).

**Table 2:**
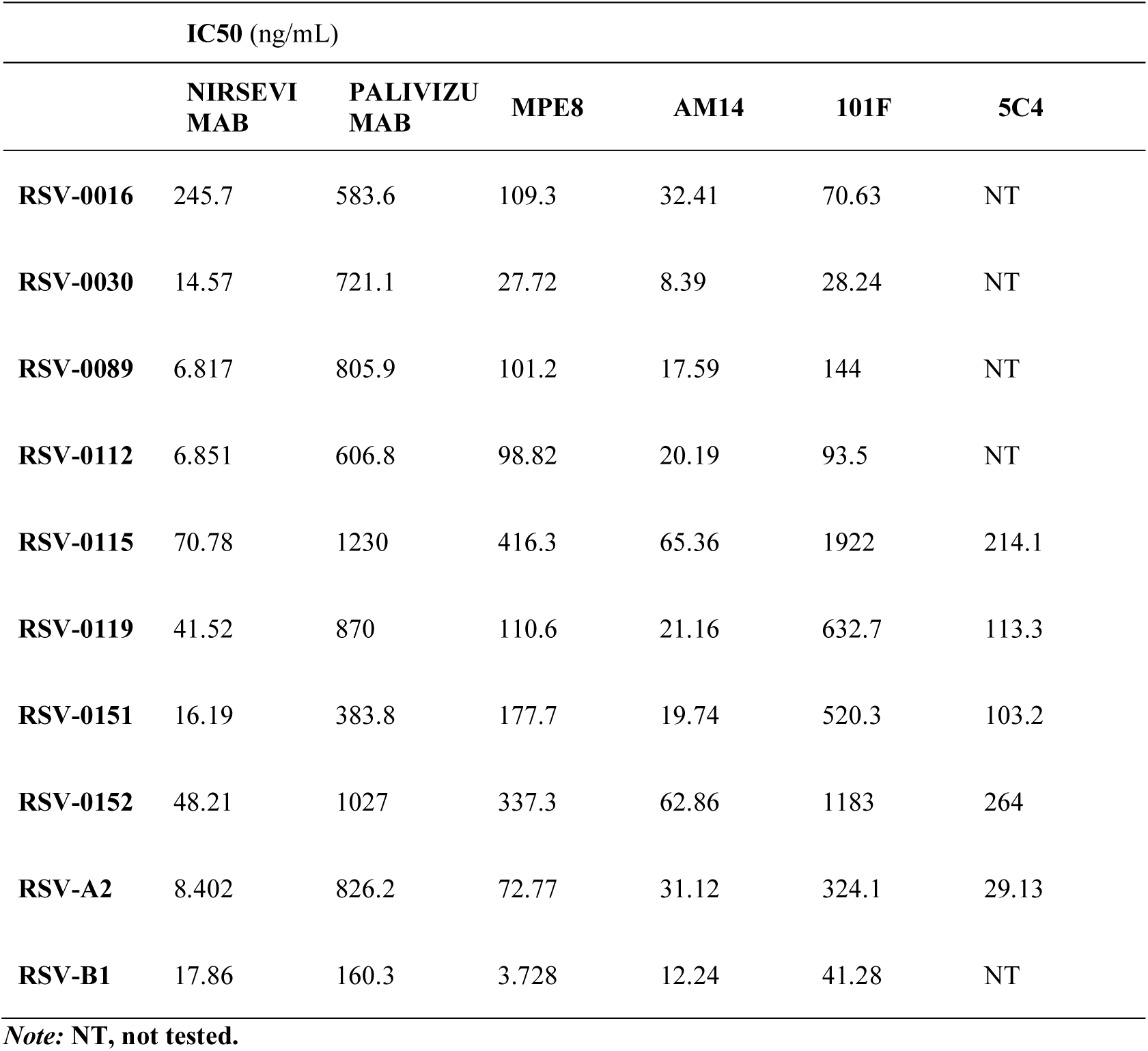
*In* vitro neutralization potency of mAbs against RSV isolates. IC50 values (ng/mL) represent the antibody concentration required to achieve 50% viral neutralization.

Antibodies with different epitope specificities displayed distinct sensitivity patterns. The prefusion-specific antibody AM14 maintained robust neutralization across most isolates, with IC₅₀ values ranging from approximately 8 to 65 ng/mL. Conversely, antibodies such as 101F and 5C4, which target defined antigenic sites on the F protein, exhibited greater variability. These combinations were not tested (NT), as 5C4 is known to specifically bind RSV-A strains at antigenic site 0, and therefore no activity against RSV-B is expected.

To investigate antigenic relationships among isolates among the panel of antibodies, we performed antigenic mapping using principal component analysis (PCA) using the IC₅₀ matrix (Figure 7). The PCA biplot revealed separation between RSV-A and RSV-B isolates, indicating subtype-specific neutralization patterns. Vectors representing the mAbs showed distinct orientations, reflecting their differential influence on the observed variance. Nirsevimab contributed strongly along PC1, consistent with its high discriminatory power across isolates, whereas 101F exerted substantial influence along PC2. In contrast, antibodies such as MPE8, AM14 and palivizumab clustered more closely, indicating more similar response profiles across the tested isolates. This analysis shows that overall antigenic differences to a panel of mAbs separate RSV subtypes, even when differences to individual mAbs are small.

**FIGURE 7:**
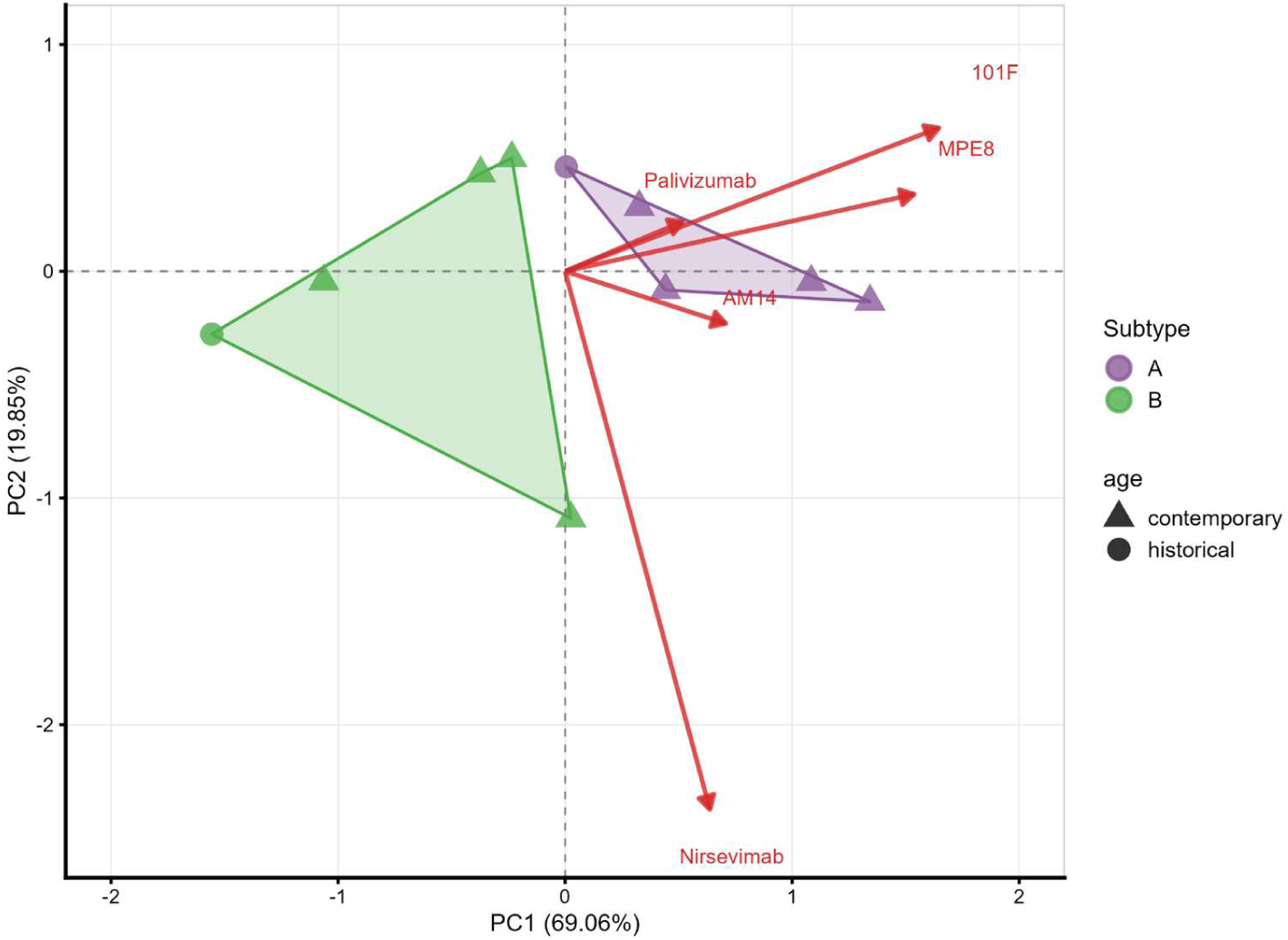
Principal component analysis (PCA) of RSV neutralization sensitivity to monoclonal antibodies. PCA was performed on IC₅₀ values from the mAb panel. Each point represents an RSV isolate, with shading indicating the convex hull of each subtype (red = RSV-A; teal = RSV-B). Arrows denote the loading vectors, reflecting the contribution and directionality of each mAb to the variance. Nirsevimab and 101F show the strongest influence along PC1 and PC2, respectively, driving separation of isolates. RSV-A and RSV-B clusters segregate distinctly, consistent with subtype-specific neutralization profiles.

### Polyclonal response using rabbit serum

We next assessed polyclonal responses. Using the FRNT assay, we assessed neutralization responses to Rabbits immunized with mRNA-1345 (RSV-A preF), mRNA encoding the RSV-B preF antigen, or with a combination of both (ratio 1:1). Neutralization titers (ID₅₀) for each group are shown in Figure 8. The RSV-A immunization regimen induced robust neutralization across multiple RSV-A strains, with particularly high titers against RSV-A119 and RSV-A115. In contrast, RSV-B immunization generated more restricted responses, with consistently lower titers against most RSV-A strains. The combined RSV-A/B immunization resulted in the broadest neutralization profile, with strong responses across both subtypes and markedly elevated titers against specific RSV-A and RSV-B strains.

**FIGURE 8.**
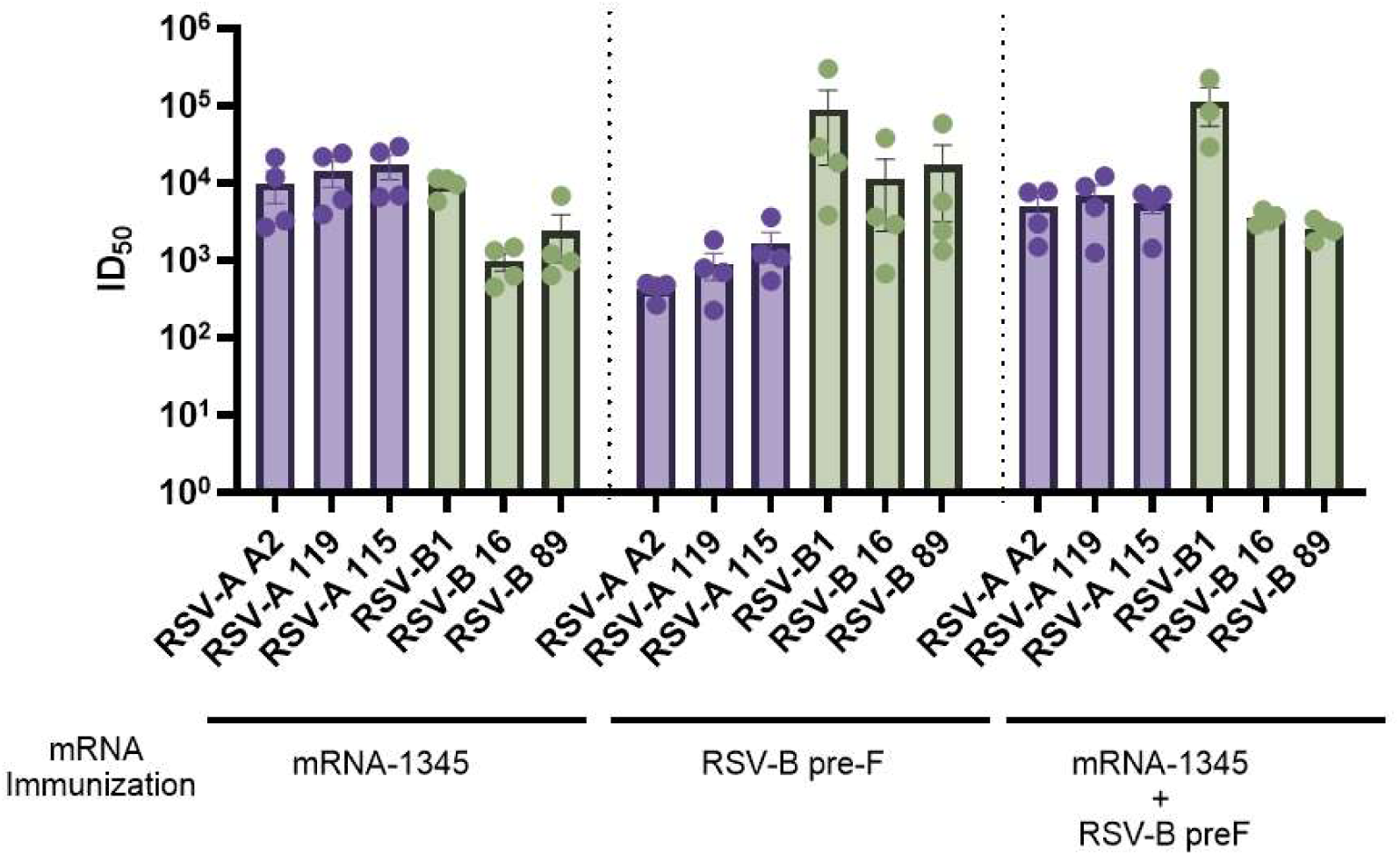
RSV neutralization titers in sera from rabbits vaccinated with mRNA-1345, RSV-B preF, or mRNA-1345 + RSV-B preF (n = 4 animals per group). ID50 neutralization titers are shown for RSV-A and RSV-B strains as indicated and Bars represent mean ID50 neutralization titers ± SEM, with individual animal values (n = 4 per group) shown as dots.

To assess global patterns among the isolates and immunogens tests, we repeated the antigenic mapping procedure used for mAbs. PCA revealed clear segregation of neutralizing antibody responses according to RSV subtype (Figure 9). Samples corresponding to RSV-A and RSV-B formed distinct clusters along PC1, indicating subtype-specific response profiles. Within the RSV-B group, the historical isolate B1 was an outlier, suggesting antigenic divergence of contemporary isolates from this commonly used historical control. Compared to immunization with RSV-A and RSV-B monovalent vaccines, responses elicited by the bivalent RSV-A/B vaccine were intermediate between the two subtype clusters, consistent with a broader and less polarized antibody profile.

**FIGURE 9.**
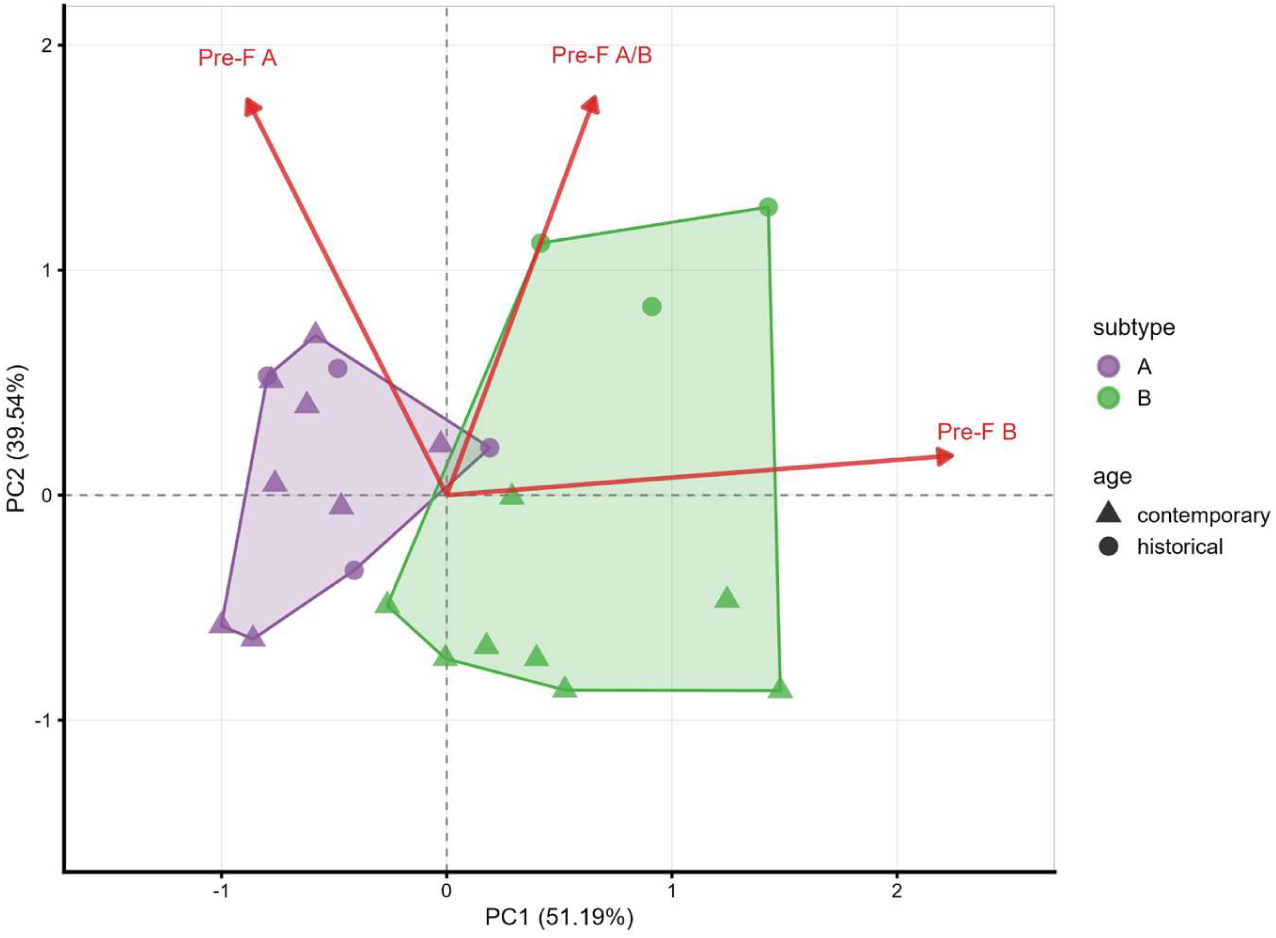
Principal component analysis (PCA) of rabbit sera responses to RSV vaccination. Individual samples are plotted along the first two principal components (PC1 = 46.43%; PC2 = 43.73%). Points are colored by vaccine subtype (A, red; B, blue) and shaped by age category (historical, circles; present, triangles). Shaded polygons denote 95% confidence ellipses for each subtype. Arrows indicate the loading vectors for variables contributing to the separation among groups.

## Discussion

In this study, we analyzed the genetic and phenotypic divergence of contemporary RSV isolates. We recovered a large number of viable isolates from both RSV-A and RSV-B positive samples and selected phylogenetically representative isolates for phenotypic analysis, including growth kinetics and antigenic mapping using a panel of mAbs and monovalent and bivalent vaccination.

Our approach to viral isolation is similar to previous studies in which clinical isolates from nasal or nasopharyngeal swabs and aspirates were successfully cultured in vitro HEp-2 or Vero, allowing generation of low-passage stocks for downstream phenotypic analysis ^27^. After isolation, we re-sequenced the viral genomes and found no nucleotide differences between the original clinical samples and the corresponding cultured isolates, indicating preservation of the original viral sequence through the isolation process. The use of low-passage, genetically stable isolates is critical, because as noted by previous studies, many published RSV data rely on long-passaged lab strains, which may accumulate cell-culture adaptive mutations that alter viral phenotype and antigenicity ^28^. Consequently, our low-passage, sequence-verified isolates provide a more physiologically relevant tool for immunological and neutralization assays.

Our findings have epidemiological significance. The isolates tested represented the genetic subtypes that were associated with the particularly severe RSV 2022-2023 season. The similarity in growth kinetics and neutralization profiles of these isolates to historical controls further supports the idea that non-viral factors were responsible for that early and severe RSV season.^22^

Genetic and phenotypic diversity also has implications for the ongoing surveillance and future development of immunoprophylaxis agents. Mutations in the mAb binding site can alter antibody efficacy. Suptavumab failed in development because specific mutations in the F protein, primarily F:L172Q and F:S173L in RSV-B strains, led to a complete loss of neutralization activity.^29^ These resistance-associated mutations were responsible for the failure of suptavumab in clinical trials and ultimately led to the discontinuation of its development ^29^. However, mutations within mAb binding sites do not always confer resistance. For example, F:I206M and F:Q209R in the nirsevimab binding site of RSV-B clinical isolates are not associated with reduced antibody sensitivity^30^. Because some amino acid changes can dramatically affect mAb potency, whereas others may be tolerated without compromising neutralization^31^, it remains critical to perform phenotypic testing on genetically variable isolates.

We evaluated a panel of well-characterized mAbs targeting defined epitopes on the RSV F protein against contemporary RSV A and B virus isolates.Simonich et al.^32^ showed that pseudoviruses expressing F proteins from both recent and historical RSV A and B strains were neutralized by nirsevimab as well as palivizumab, clesrovimab, AM14, and MPE8. Consistent with these findings, our results indicated that the major circulating subtypes of RSV A and B remain susceptibility to all mAbs tested, RSV A strains required slighlty higher antibody concentrations to achieve neutralization compared to RSV B strains. Other studies have similarly reported subtype-dependent differences, showing that RSV-B isolates are more easily neutralized by mAbs than RSV-A ^27,30^.

One isolate, RSV-B16 isolate showed an IC_50_ to nirsevimab approximately 13-fold higher than the RSV-B1 control and all other viruses tested, including RSV-A strains. Notably, this difference occurred in the absence of a mutation in the nirsevimab binding site. Stobbelaar et al. ^30^ also reported that amino-acid variation within the F protein does not always correlate with differences in susceptibility to mAb neutralization, suggesting that additional viral factors may modulate antibody activity. Indeed, recent work has shown that co-expression of the attachment glycoprotein G can alter the conformation of F and significantly enhance the binding and neutralization potency of anti-F mAbs, indicating that F-only recombinant viruses may not fully recapitulate the antigenic context of native virions ^33^. Taken together, these data support our observation that none of the clinical isolates in this study carry mutations within known mAb binding sites that could explain the reduced susceptibility. Instead, factors outside the F sequence itself, such as epistatic effects, virion architecture, or interactions between F and other viral proteins, may contribute to the resistance of phenotype observed in a subset of isolates.

In rabbit immunization studies, contemporary RSV-B isolates displayed a distinct neutralization profile compared with the laboratory strain RSV-B1. In contrast, sera challenged with RSV-A strains showed highly consistent neutralization patterns, similar to the historical reference isolate A1. A similar strain-dependent effect was described in the evaluation of RSVPreF3 immunogenicity ^34^ where non-adjuvanted RSVPreF3, based on the RSV-A2 sequence, boosted neutralizing titers more effectively against RSV-A and RSV-B laboratory strains than against contemporary RSV-B viruses, as well as studies conducted by Hause et al^35^. showing that differences between historical and contemporary RSV-A sequences can lead to reduced efficacy against RSV-B subtype viruses. RSV B1 and A2 are commonly used as prototype sequences in vaccines and diagnostic assays (e.g. ^36–39^). Our findings suggest that RSV-B1 is antigenically distinct from contemporary RSV-B viruses and findings from RSV B1 may not generalize to circulating RSV-B viruses.

The genetic and phenotypic differences among contemporary RSV are effectively bimodal and summarized by diverged, co-circulating subtypes. The bivalent pre-F vaccine formulation tested in our study elicited highly comparable neutralizing responses across all viruses evaluated, both laboratory strains and contemporary clinical isolates, supporting the idea that inclusion of both RSV-A and RSV-B antigens can mitigate subgroup-specific or strain-specific differences in antibody recognition. Our results also align with previous studies, including Nuttens *et al*, ^10^, which indicate that a bivalent RSV A/RSV B vaccine may provide broader and more durable protection across subgroups compared with monovalent vaccines or mAbs. This, bivalent immunization may be an effective strategy to address the effectively bimodal genetic and phenotypic diversity of contemporary RSV isolates.

## Methods

### Cells and Viruses

Hep2 (CCL-23, ATCC) and A549 cell lines (CCL-185, ATCC) were cultivated in Dulbecco’s Modified Eagle Medium (DMEM) supplemented with 10% heat-inactivated fetal bovine serum (FBS), 100 IU/mL penicillin and 100μg/mL streptomycin (all reagents from Gibco). All cells were kept at 37°C in a humidified, 5% CO2-controlled atmosphere and were passaged twice a week.

RSV-A reference strains A2 (NR-44233) and RSV-B reference strain B1 (NR-56243) were obtained from BEI resources. The viruses were cultivated in HEp-2 cells until cytopathic effect (CPE) was visible throughout the flask. The virus was collected and quantified in a conventional plaque assay.

### Specimens

Nasal/nasopharyngeal swab specimens were collected from patients who tested positive for RSV by RT-PCR at Massachusetts General Hospital between November 2022 and March 2024. A total of 137 specimens were subtyped using next-generation sequencing analysis.

### Phylogenetic Analysis

RSV-A and RSV-B genome sequences from clinical isolates collected in Massachusetts between seasons 2022 to 2024 were combined with publicly available reference sequences from NCBI database. For RSV-A, we included 394 historical sequences (1977-2019) and 487 contemporary sequences (2020-2024). For RSV-B, we included 204 reference sequences spanning 1970-2024. All sequences were classified into genotypes using Nextclade v3 (https://clades.nextstrain.org/) with RSV-A and RSV-B datasets. Sequences with quality control status of “good” were retained for phylogenetic analysis.

Multiple sequence alignments were generated using MAFFT v7 (https://mafft.cbrc.jp/alignment/software/) as implemented in Augur (https://docs.nextstrain.org/projects/augur/)). Genome sequences with Nextclade quality control status of ‘good’ and genomic coverage >99% were retained for analysis, resulting in 55 RSV-A and 46 RSV-B high-quality sequences. Whole genome sequences that did not meet these quality criteria were excluded from analysis. Maximum likelihood phylogenetic trees were inferred using IQ-TREE with automatic model selection. Temporal calibration was performed using TreeTime with a coalescent model (--coalescent opt) to estimate divergence times and evolutionary rates. Trees were visualized using Auspice v2 (https://docs.nextstrain.org/projects/auspice/), with sequences colored by Nextclade-assigned clade and sample source.

### Genetic Divergence Analysis

To quantify intra-group genetic diversity, we calculated nucleotide divergence of each clinical isolate relative to its respective subtype consensus sequence. We aligned genomes using MAFFT and extracted F and G regions based on GenBank annotations. For each subtype (RSV-A and RSV-B), we generated a consensus sequence from all quality-filtered clinical isolates, then calculated divergence of each sample as the percentage of nucleotide differences at non-gap positions relative to this consensus. We filtered sequences for quality (>75% length, <5% N content), with filtering applied independently to each gene alignment, resulting in varying sample sizes between genes due to differences in sequencing quality across genomic regions (Gene F: RSV-A n=53, RSV-B n=45; Gene G: RSV-A n=63, RSV-B n=42). For group comparisons, we used Mann-Whitney U tests due to non-normal data distribution (Shapiro-Wilk p < 0.05), with Bonferroni correction for multiple comparisons (α = 0.025).

For phylogenetic visualization, we identified nucleotide and amino acid mutations relative to reference sequences RSV-A2 (KT992094.1) and RSV-B1 (AF013254.1). Whole genomes were aligned using MAFFT v7, and F gene (RSV-A: nt 5662–7386; RSV-B: nt 5666–7390) and G gene (RSV-A: nt 4689–5585; RSV-B: nt 4690–5589) regions were extracted based on GenBank annotations. We excluded fixed mutations (present in 100% of samples) from plots to focus on variable sites. For amino acid analysis of the F protein, nucleotide alignments were translated and substitutions identified relative to the reference sequences. Epitope regions targeted by clinically relevant mAbs (nirsevimab, palivizumab, 5C4, MPE8, 101F, and AM14) were annotated based on published structural and mutagenesis studies. In all alignments, samples were ordered by phylogenetic relationships determined using Nextclade v3, to highlight clade-specific mutational patterns.

We performed bioinformatic analyses using Nextstrain/Augur v24, Nextclade v3, MAFFT v7, and custom Python scripts. For some analyses, Claude (Anthropic) provided draft code and guidance on pipeline implementation and data visualization, with draft code reviewed for accuracy by the investigators.

### Monoclonal Antibodies

We obtained sequences of the heavy and light chains for mAbs AM14, MPE8, D25 (nirsevimab), palivizumab, 101F and 5C4 from referenced publications and structures in the Protein Data Bank. We synthesized DNA encoding antibody heavy and light variable regions, codon-optimized for human expression. AM14, MPE8, D25, Palivizumab, and 101F heavy and light variable regions were subcloned into expression plasmids containing in-frame human constant domains (IgG1 for heavy chain and kappa for light chain). 5C4 heavy and light variable regions were subcloned into expression plasmids containing in-frame mouse constant domains (IgG2a for heavy chain and kappa for light chain). We expressed antibodies by transient co-transfection of heavy and light chain plasmids into ExpiCHO cells in suspension at 37°C for 3-5 days, passed cell supernatents over Protein A agarose and eluted bound antibodies with 1 M Tris-HCl pH 8.0, with further purification by size-exclusion chromatography (GenScript).

### Virus Isolation from Clinical Samples

We stored nasal/nasopharyngeal swabs at −80 °C before processing in cell culture. The nasopharyngeal swabs were inoculated to Hep-2 cells cultured in Dulbecco’s modified Eagle’s medium supplemented with 10 % fetal bovine serum. We assessed CPE of inoculated Hep-2 cells every day for seven days. CPE-negative samples on the seventh day after inoculation were blindly inoculated for two consecutive passages, and CPE-positive samples were recorded and stored at −80 ◦C for future use.

Once CPE was observed, we confirmed viral isolation by immunostaining and next-generation sequencing (MiniSeq, Illumina, San Diego, CA, USA). To prepare viral stocks, we infected HEp-2 cells at a multiplicity of infection (MOI), representing the ratio of infectious viral particles to target cells, of 0.01 and harvested after 72 h. We titered stocks using plaque assays in Hep-2 cells.

### Immunostaining

We fixed plates with 3.7% formaldehyde for at least 10 min at room temperature, followed by washing with phosphate-buffered saline (PBS). Cells were then permeabilized with PBS containing 0.01% Triton X-100 for 15 min and washed again with PBS. We then incubated cells with an RSV polyclonal antibody conjugated to FITC (Invitrogen, PA1-73017) at a 1:500 dilution for 3 h at room temperature under gentle rocking and measured fluorescence using a Cytation 5 cell imaging multi-mode reader (BioTek Instruments, Winooski, VT, USA).

### Plaque assay

We performed plaque assays using HEp-2 cells to quantify infectious virions. Briefly, we infected Hep-2 cells with 10-fold serial dilutions of virus isolate at 37°C in 5% CO2 for 1 h. We then removed the inoculum and added 3 mL of an overlay medium (composed of 0.4% Carboxymethyl cellulose with 199 media (Cat. no. 21157029, Thermofisher), containing 10% FBS). We fixed infected cells with 3.7% formaldehyde for at least 3h and then stained with crystal violet to count the number of plaques after 7 days. We determined viral titers by counting plaque-forming units (PFU).

### Neutralization Assays

#### Fluorescent Focus Reduction Neutralization Assay

Eight serial 2-fold dilutions of mAbs were made in 40ul DMEM in 96 well microtiter plates. Forty microliters of virus containing ∼ 100 pfu of RSV A or B strain were added to the mAbs dilutions and allowed to incubate for 1h at 37°C in 5% CO2. Positive control wells of virus without mAbs and negative control wells without virus or mAbs were included on each plate. Then, 50 ul the mAbs-virus mixtures were added to the Hep-2 plates incubated at 37°C in 5% CO_2_ for 1h, the inoculum was removed, 100 ul overlay medium was applied, and the plates were again incubated at 37°C in 5 % CO_2_ for 28h. The plates were fixed with 4% formaldehyde for 10 min. Subsequently, the cells were permeabilized and blocked for 10 min with PBS containing 0.1% TritonX-100, followed by incubation with an antibody targeting the RSV (Invitrogen, PA1-73017) for 30 min in room temperature. After the final wash, images were captured under an 4x objective lens using Cytation 5. Inhibitory concentration leading to 50% viral neutralization (IC50) was calculated based on 8-point dilution series using a non-linear curve fitting in GraphPad Prism 10.0 software (GraphPad Software, La Jolla, CA, USA)

### Plaque Reduction Neutralization Assay

We mixed F-specific mAbs with an equal amount of virus suspension containing 150 plaque-forming units at 37°C for 1 h. We inoculated six-well plates containing HEp-2 cells monolayers with 80% confluence with virus/mAb solution and incubated at 37°C for 1 h. Subsequently, we removed the inoculum and added 3 mL of an overlay medium. After 7 days, we fixed plates with 3.7% formaldehyde for 6h, stained with crystal violet, and counted the plaques.

### Viral Replication Kinetics

We seeded Hep-2 and A549 at a density of 5 × 10^5^ cells per well in 6-well plates the day before infection. We diluted the virus in DMEM to achieve an MOI of 0.01 and incubated this mixture with target cells for 1 h at 37°C. After 1 h, we removed the inoculum, washed cells with PBS twice, replaced the media with fresh media containing 10% FBS, and incubated the mixture at 37°C in 5 % CO2. We collected supernatants daily up to day 4 post-infection and stored them at −80 ◦C. We assessed replication kinetics by plaque assays.

### Rabbit serum

We immunized 12 female New Zealand White rabbits, 30 weeks old, in three treatment groups (n=4 rabbits per group). We injected animals with 20 ug of mRNA-1345 (RSV A preF), 20 ug of mRNA encoding RSV B preF, or 10 ug mRNA-1345 plus 10ug of RSV B preF formulation via the intramuscular route on Days 0, 21, and 42. We collected blood on Days 0 (pre-bleed), 36, and 63. We humanely euthanized animals following terminal blood collection on Day 63. The study complied with applicable sections of the current versions of the following regulations and guidances: 1) Animal Welfare Act Regulations (9 CFR); 2) U.S. Public Health Service Office of Laboratory Animal Welfare (OLAW) Policy on Humane Care and Use of Laboratory Animals; 3) Guide for the Care and Use of Laboratory Animals (Institute of Laboratory Animal Resources, Commission on Life Sciences, National Research Council, 2011); and 4) Applicable Noble Life Sciences’ SOPs.

### Statistical analysis

Statistical analyses were performed using GraphPad Prism X. IC₅₀ values obtained from neutralization assays with mAbs were compared between RSV-A and RSV-B using a two-tailed Mann–Whitney U test (non-parametric). A p value < 0.05 was considered statistically significant.

To explore patterns in neutralizing antibody responses, we used principal component analysis (PCA)^25^ to carry out antigenic cartography ^26^. We displayed the first two components to visualize responses by RSV subtype.

### Funding

This study was funded by Moderna Tx and the Massachusetts Consortium on Pathogen Readiness (GRO245623). J.L. was also supported by an NIH grant (R37 AI147868).

## Acknowledgments

The following reagents were obtained through BEI Resources, NIAID, NIH: Human Respiratory Syncytial Virus, A2, NR-12149 and Human Respiratory Syncytial Virus, B1, NR-56243.

## Data Availability

The authors confirm that the data supporting the findings of this study are available within the article and its supplemental material.

